# Using population-level whole-genome sequencing to profile colistin resistance evolution dynamics in diverse clinical *Pseudomonas aeruginosa* lineages

**DOI:** 10.64898/2026.09.22.753656

**Authors:** Kellie R. Strickland, Miriana L. Littlejohn, Erin P. Price, Derek S. Sarovich

## Abstract

Colistin (CST) is a critical last-line antibiotic for treating multidrug-resistant *Pseudomonas aeruginosa* infections, especially in chronic respiratory disease. Despite its clinical importance, the *P. aeruginosa* CST resistome remains incompletely characterised, with most studies focusing on the PAO1 prototypic strain. Here, we used experimental evolution to apply stepwise CST selective pressure to nine *P. aeruginosa* lineages representing different clinical presentations and diverse genetic backgrounds. Population-level whole-genome sequencing of broth cultures exposed to 2-fold increasing CST (0.0625 to 512 μg/mL) was undertaken at multiple concentrations. We identified several known and novel mutational drivers of low- and high-level CST resistance, alongside associated compensatory mutations at allele frequencies (AFs) as low as 5%. Despite diverse clinical and genetic backgrounds, all lineages convergently evolved mutations within the two-component system *pmrAB,* with most accompanied by *phoPQ* variants. When canonical TCS remodelling plateaued, populations acquired novel secondary driver mutations, most notably in the lipid A-modifying gene, *lpxO2*. In addition to AMR driver mutations, strains evolved complex adaptive strategies to mitigate severe CST-induced oxidative and metabolic stress. Adaptations included the emergence of hypermutators and alterations to targeted DNA and protein repair networks. Furthermore, we observed convergent regulatory rewiring of the cyclic-di-GMP network and central metabolism pathways, likely acting as compensatory mechanisms to promote biofilm formation and offset the fitness costs of CST resistance. Using a mixture-aware population genomics approach, we captured intense clonal interference and transient tolerance mutations that failed to reach fixation, illustrating dynamic evolutionary complexities that would be missed with traditional end-point analysis methods.

## Introduction

*Pseudomonas aeruginosa* is a serious global public health concern due to its environmental ubiquity, pathogenicity, persistence, high antimicrobial resistance (AMR) rates, and association with increased morbidity across a spectrum of diseases^1,2^. This Gram-negative pathogen is particularly burdensome for people with chronic respiratory conditions such as cystic fibrosis (CF), bronchiectasis, and chronic obstructive pulmonary disease (COPD), where long-term colonisation is common. This persistence is problematic and directly associated with lung function decline, poorer quality of life, increased likelihood of exacerbations, and increased mortality risk^3–5^.

First discovered in 1947, CST (polymyxin E) fell out of favour in the 1970s due to high nephro- and neuro-toxicity rates when delivered intravenously^6^. However, CST is experiencing a renaissance in the AMR era, particularly in nebulised or topical form^6^. Nebulisation achieves high localised doses, effectively targeting respiratory infections whilst reducing systemic toxicity risks^6^. Due to its low historical use, older surveillance studies report low CST resistance rates in *P. aeruginosa* (∼1% global prevalence); however, a recent meta-analysis found a slow but insidious increase, currently at 5% overall, and 7% in CF-derived isolates^7^. Historically, Australia and New Zealand have reported the lowest CST resistance rates, largely due to their geographic isolation, stringent biosecurity measures, no historical CST use in agriculture, and its strict reservation as a last-line therapeutic. Although rare in this region, CST-resistant isolates have been identified in people with CF^8^ or ocular infections^9^; however, no CST-resistant strains have been identified in wild animals, livestock, or domestic animals^10^. Importantly, no *mcr-1*-harbouring *P. aeruginosa* strains have been reported in Australia or New Zealand.

The globally low CST resistance rate is also attributable to its predominantly chromosomal mutation basis, which typically incurs severe fitness cost^11,12^. Despite its rarity, CST resistance in *P. aeruginosa* threatens to be catastrophic, with a small case study reporting a 30-day mortality rate of 87.5% for eight patients with CST- and carbapenem-resistant infections^13^. Although currently uncommon, the increasing global use of nebulised CST to treat carbapenem-resistant *P. aeruginosa*, *Acinetobacter baumanii*, and *Klebsiella pneumoniae* lung infections is now risking the future effectiveness of this ‘last-line’ antibiotic^14–16^. Additionally, the emergence and global dissemination of plasmid-borne *mcr-1* CST resistance genes further jeopardises the future of successful CST treatment^17–19^.

Belonging to the polymyxins, CST is an attractive anti-pseudomonal antibiotic due to its relatively low risk of cross resistance, with strains typically remaining susceptible to CST even when other antibiotic options are exhausted^20,21^. Mechanistically, CST solubilises the bacterial cell membrane by binding to the lipid A component of lipopolysaccharide (LPS). Electrostatic interactions then displace stabilising cations within the LPS, allowing CST to insert its hydrophobic tail into the outer cell membrane, promote its own uptake, destabilise the inner membrane, and ultimately cause cell lysis and death^21–23^. The main resistance mechanism in *P. aeruginosa* occurs via LPS modification, preventing CST electrostatic interactions with the negatively-charged (anionic) outer cell membrane^11,12,20,21,24–27^. *P. aeruginosa* can reduce its lipid A anionic charge by upregulation of the *arn* operon, generally through activation of one of its two-component systems (TCSs), which subsequently adds the cationic sugar molecule, 4-amino-4-deoxy-L-arabinose (L-Ara4N) to lipid A. Single mutations in these TCSs are frequently sufficient to push the CST minimum inhibitory concentration (MIC) above standard clinical breakpoints; however, high-level CST resistance often requires multiple, synergistic mutations^11,21^.

Despite this mechanistic knowledge, critical gaps remain regarding the evolutionary dynamics of CST resistance in *P. aeruginosa*. Most prior studies have relied on the laboratory-domesticated strains PAO1 and PA14^28–32^, which fails to capture the phenotypic diversity observed in contemporaneous clinical isolates, such as alginate overproduction (mucoidy) and hypermutation. This knowledge gap is particularly relevant in the Australian context, which has genetically distinct *P. aeruginosa* populations compared with more comprehensively studied European and American lineages^33,34^., Consequently, it remains unknown whether these different genotypes may evolve CST resistance via alternative genetic variants, mechanisms, or pathways than those characterised in Euro-American clades. Moreover, prior experimental models often fail to reach the high resistance levels seen in naturally evolved clinical isolates; for example, strains derived from CF cohorts can exhibit CST MICs >256 µg/mL. As research has heavily prioritised CF-derived strains, the *P. aeruginosa* lineages more commonly associated with COPD and bronchiectasis have been largely overlooked. Finally, although bioinformatic approaches for predicting phenotypic CST resistance from whole-genome sequences have been implemented, they remain underdeveloped and are hindered by high false-negative prediction rates^33^.

To address these knowledge gaps, we subjected nine contemporary clinical *P. aeruginosa* strains from CF, bronchiectasis, COPD, and acute bloodstream infections to experimental evolution under increasing CST pressure. Strains were chosen to represent common contemporary circulating genotypes in Australia and exhibited a spectrum of susceptibilities to anti-pseudomonal antibiotics, thus representing the ‘real-world’ clinical setting in our region. Next, we performed deep-sequencing of mixed *P. aeruginosa* populations across increasing CST MICs to identify drivers of CST resistance and associated compensatory mutations, and to explore AMR variant dynamics.

## Methods

### Ethical approval

Ethics approvals were obtained from Metro North Human Research Ethics Committee (HREC) (HREC/13/QPCH/127^35^, HREC/2019/QPCH/48013, HREC/2022/MNHB/87992, and HREC/18/QPCH/110^36^), the Royal Brisbane and Women’s Hospital HREC^37^, and the Mater Hospital HREC (HREC/MML/89908). Site-specific approvals were subsequently obtained from the relevant Queensland hospitals (Sunshine Coast University Hospital, The Prince Charles Hospital, Royal Brisbane and Women’s Hospital, Mater Hospital Brisbane). All participants provided written consent, except for the bloodstream and non-CF bronchiectasis isolates, where a waiver of informed consent was granted due to the low to negligible risk associated with these studies^37^.

### P. aeruginosa strains

Nine Australian clinical *P. aeruginosa* strains (bronchiectasis, *n*=2; CF, *n*=1; COPD, *n*=3; and acute bloodstream infection, *n*=3), all susceptible to CST (MIC ≤1μg/mL), were examined in this study (Table 1). Multilocus sequence types (STs) were chosen to represent the most common genotypes found in Australia, including the high-risk international clone, ST274^38^. A variety of antibiotic susceptibility profiles were included to cover the AMR spectrum, and two strains exhibited mucoidy. In addition, PAO1 (LMG 12228; Belgian Coordinated Collections of Microorganisms, Ghent University, Gent, Belgium), which was isolated from an infected wound in Melbourne in 1954^39^, was included as a historical Australian clinical control.

**Table 1.** *Pseudomonas aeruginosa* strains used in this study.

| Strain | Disease | ST | Mucoidy | AMR variant/s* | CST MIC (µg/mL) | AMR profile |
| --- | --- | --- | --- | --- | --- | --- |
| SCHI0021.S.7 | CF | 801 | Yes | AmpD P42fs, FusA1 S459F, GyrA D87H, RplB G138S | 0.5 | AMK<sup>r</sup>, TOB<sup>r</sup>, FEP<sup>r</sup>, CIP<sup>r</sup> |
| SCHI0033.S.8 | BSI | 649 | No | DacB G427D | <0.25 | PIP<sup>r</sup>, TZP<sup>r</sup>, CAZ<sup>r</sup>, FEP<sup>r</sup> |
| SCHI0033.S.19 | BSI | 235 | No | GyrA T83I, ParC S87L, ParS A149T, PA0027 L34N, ArgS D184G, bla<sub>VEB-1</sub>, aph (3')-VI | 0.25 | AMK<sup>r</sup>, TOB<sup>r</sup>, PIP<sup>r</sup>, CAZ<sup>r</sup>, FEP<sup>r</sup>, CIP<sup>r</sup>, IPM<sup>i</sup> |
| SCHI0057.S.9 | BSI | 244 | No | Nil | 0.5 | Nil |
| SCHI0058.S.1 | COPD | 267 | No | PA0908 loss | 0.5 | Nil |
| SCHI0064.S.1 | COPD | 235 | No | PA0027 K34N | 1.0 | Nil |
| SCHI0109.S.2 | COPD | 385 | No | AmpD Val10Gly, GyrA Asp87Tyr, MexT loss | 1.5 | AMK<sup>r</sup> |
| SCHI0181.S.2 | BE | 395 | No | Nil | 0.5 | Nil |
| SCHI0181.S.32 | BE | 274 | Yes | AmpR D135N | 0.5 | FEP<sup>r</sup>, CIP<sup>r</sup> |
| PAO1 | Wound | 549 | No | Nil | 0.5 | Nil |

Abbreviations: AMK, amikacin; AMR, antimicrobial resistance; BE, bronchiectasis; BSI, bloodstream infection; CAZ, ceftazidime; CF, cystic fibrosis; CIP, ciprofloxacin; CST, colistin; FEP, cefepime; IPM, imipenem; MEM, meropenem; MIC, minimum inhibitory concentration; PIP, piperacillin; TOB, tobramycin; TZP, piperacillin/tazobactam; ST, multilocus sequence type; TOB, tobramycin; r, resistant; i, intermediate/increased dosage
\*Identified using ARDaP's *P. aeruginosa* database (v1.0)<sup>33</sup>.

**Table 2.**
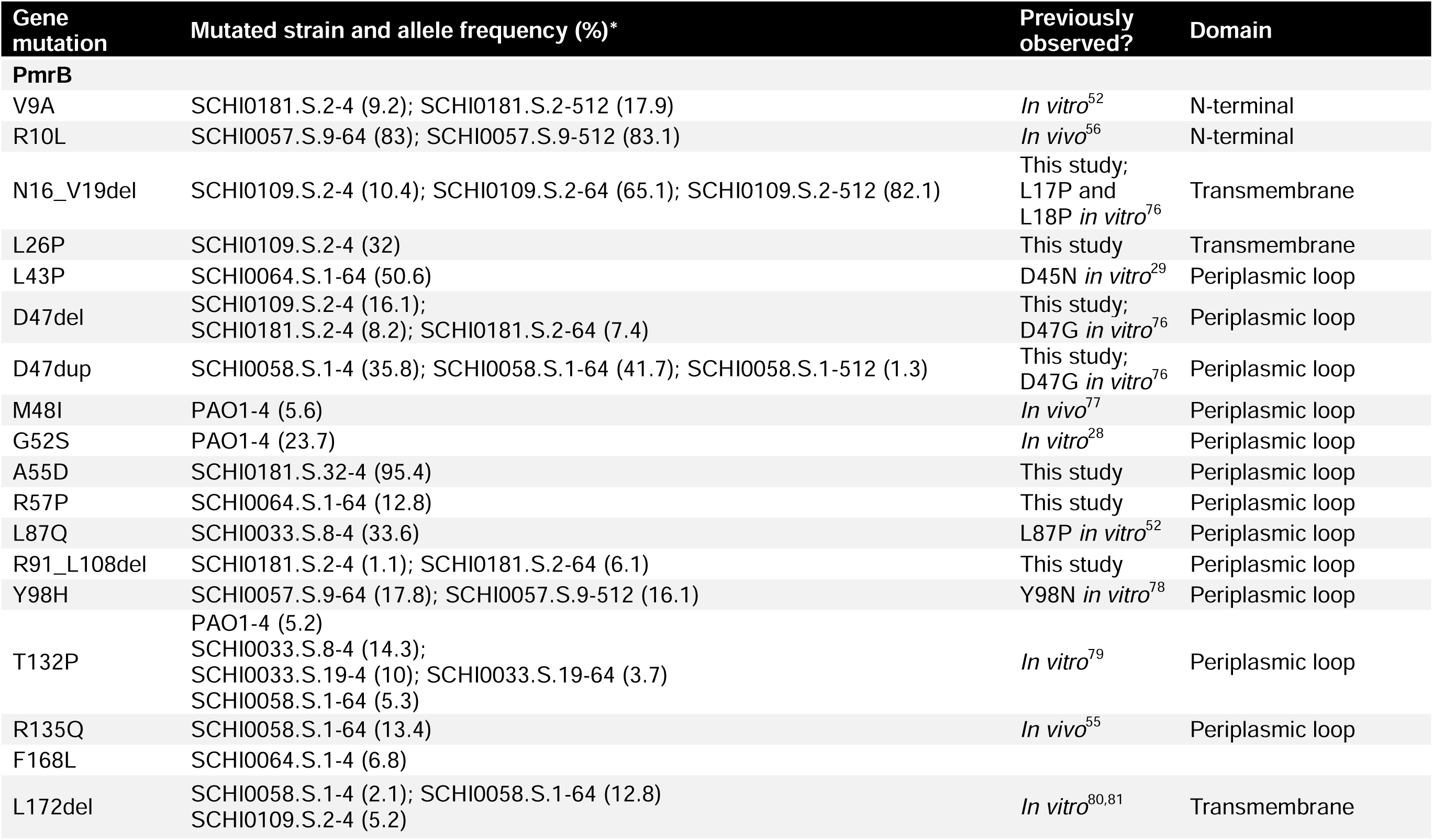
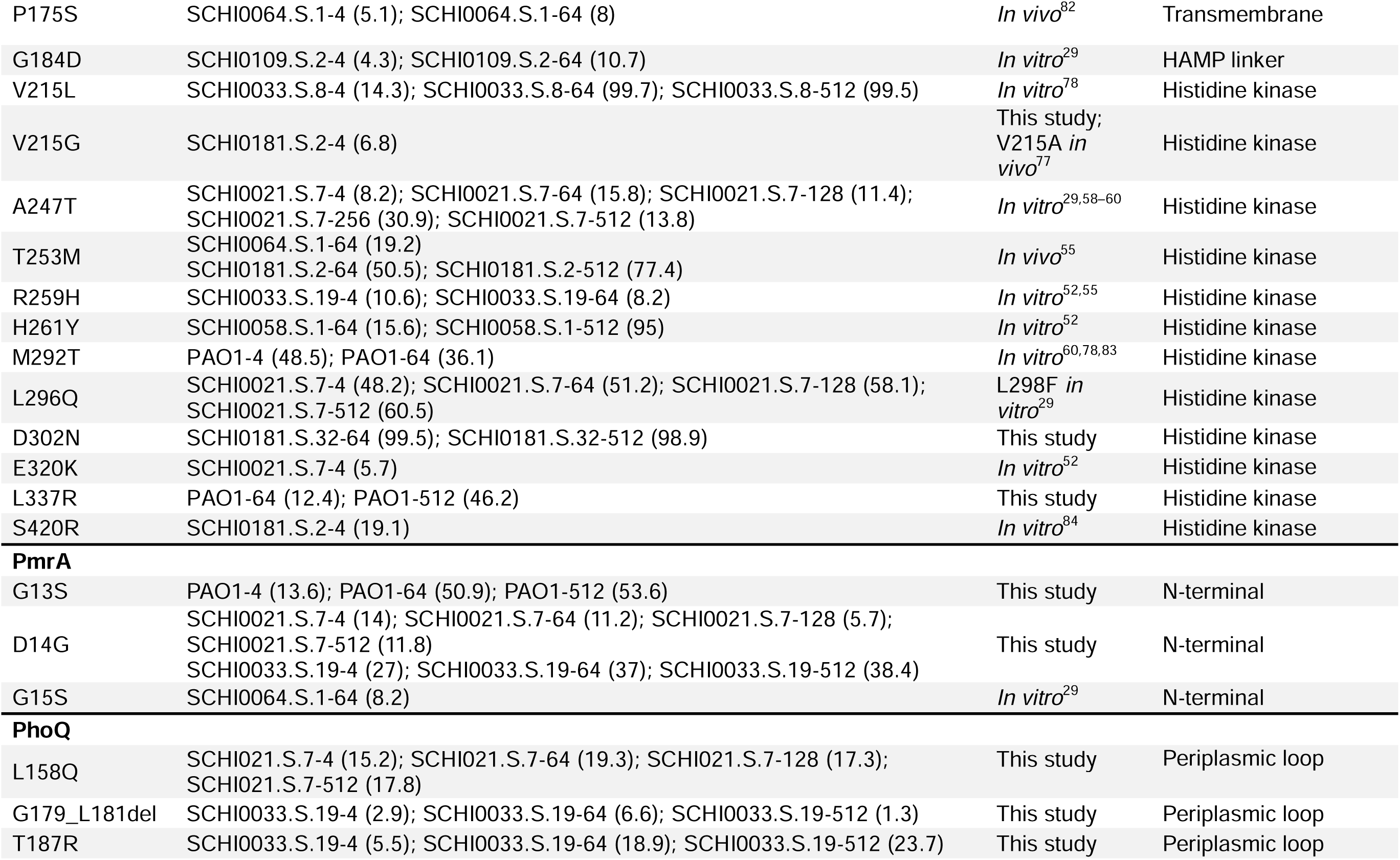
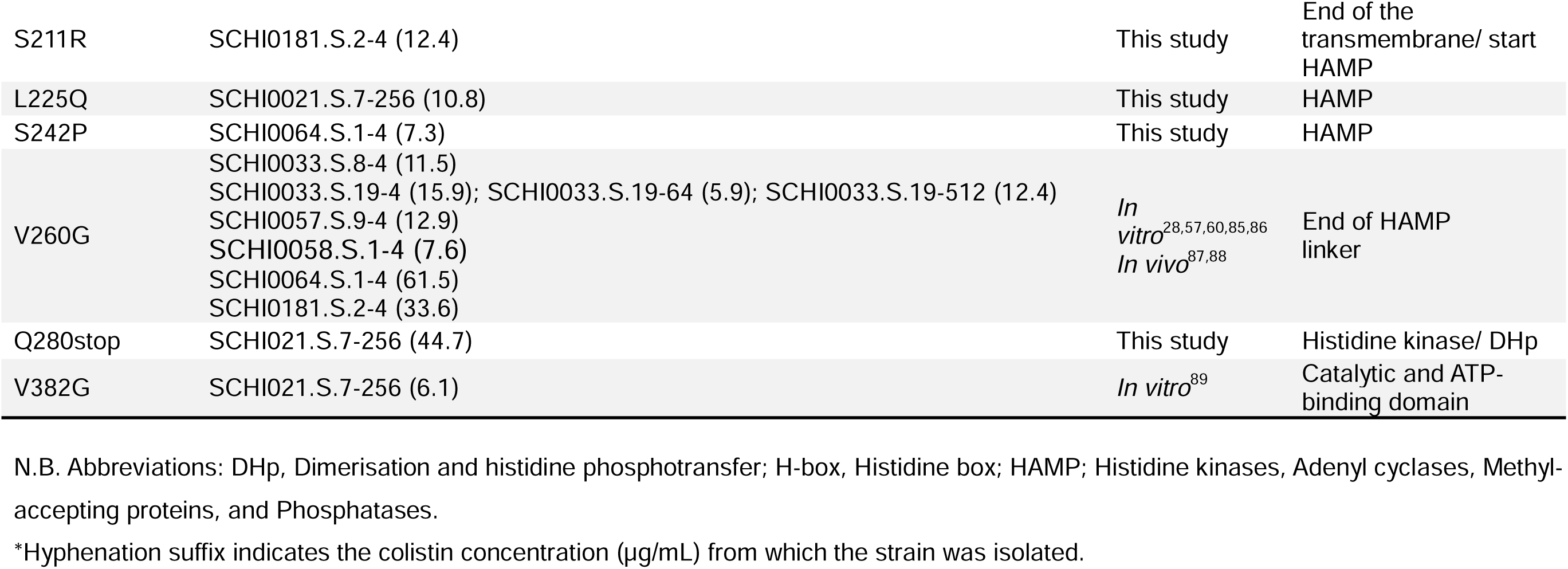
Colistin resistance driver mutations identified across ten *Pseudomonas aeruginosa* lineages exposed to increasing colistin concentrations.

### Antimicrobial susceptibility testing

Antibiotic susceptibility profiles were determined against eleven antipseudomonal antibiotics as previously described^33,40,41^, using the following disc concentrations: amikacin (AMK; 30 mg), tobramycin (TOB; 10 mg), imipenem (IPM; 10 mg), meropenem (MEM; 10 mg), cefepime (CPM; 30 mg), ceftazidime (CAZ; 30 mg), ceftolozane/tazobactam (C/T; 30/10 mg), ciprofloxacin (CIP; 5 mg), piperacillin/tazobactam (TZP; 100/10 mg), and aztreonam (ATM; 30 mg). As per EUCAST v16.0 2026 guidelines, CST MICs were determined using broth microdilution. PAO1 was included in all antimicrobial sensitivity testing as the antimicrobial-sensitive control.

### Serial passage in increasing CST concentrations

Purified *P. aeruginosa* starting inocula, equivalent to a 0.5 McFarland standard (∼1.5 × 10^8^ colony-forming units/mL), were obtained from fresh overnight cultures aerobically grown on Luria-Bertani agar at 37°C. 50 µL of *P. aeruginosa* inoculum was added to the first CST concentration (0.0625 µg/mL lyophilised CST sulphate salt, ≥19,000 IU/mg [Merck Life Science, Bayswater, VIC, Australia] suspended in 5 mL cation-adjusted Mueller-Hinton II broth [Edwards Group, Murrarie, Qld, Australia]) in a sterile 15 mL Falcon tube, to give a final bacterial concentration of ∼1 × 10^6^ cfu/mL. All *P. aeruginosa* strains, including PAO1 and the broth-only control, were incubated at 37°C for 24 h with shaking at 230 rpm using a benchtop orbital shaking incubator (Ratek Instruments [model OM11], Boronia, VIC, Australia). Where turbidity was visible after 24 h incubation, cultures were passaged by transferring 50 µL of culture to the next CST concentration in 2-fold increments up to 512 µg/mL. Where visible turbidity was not observed at 24 h, the culture was left for a further 24 h. This procedure was repeated until a concentration of 512 µg/mL was reached, or the strain became extinct (i.e. no visible turbidity after 48 h). To avoid CST degradation, all preloaded CST Falcon tubes were kept refrigerated at 4°C and in the dark until required ^42,43^. Growth at every concentration timepoint was retained by adding 200 µL of broth culture to 1mL sterile Luria-Bertani broth containing 20% glycerol, followed by storage at -80°C.

### DNA extraction and whole-genome sequencing

Total genomic DNA was extracted from all *P. aeruginosa* isolates (parental strains and experimental evolution lineages) transferring a 200μL of cultured cells into 200μL lysis buffer (20 mM Tris-HCl pH 8.0, 2 mM sodium EDTA, and 1.2% Triton X-100 [Sigma-Aldrich]), followed by bead beating in DNase-free 2mL O-ring tubes (SSIBio, Lod, CA, USA) containing ∼100μL of 0.1 and 0.5mm zirconia beads (Daintree Scientific, St Helens, TAS, Australia) using a Precellys Evolution Touch Homogeniser (Bertin Technologies, Montigny-le-Bretonneux, France) at 2 x 7500rpm for 60 sec (with a 30 sec pause between cycles). DNA was purified using the ‘Gram-positive Bacteria’ protocol of the DNeasy Blood and Tissue kit (QIAGEN, Clayton, VIC, Australia), commencing at the Buffer AL and Proteinase K addition step. Parental strains were extracted with an identical protocol but starting with ∼10ul loop of culture grown on Luria-Bertani agar. To prevent bottlenecking, experimental evolution lineages were DNA-extracted directly from broth cultures. Illumina 150bp paired-end whole-genome sequencing was performed at Macrogen (Geumcheon-gu, Seoul, Korea) or AZENTA (Suzhou, China) to achieve ∼100x coverage. For the nine parental strains, Oxford Nanopore Technologies (ONT; Oxford, UK) reads were also generated using a MinION Mk1D sequencer and R10.4.1 MinION FLO-MIN114 Flow Cells; basecalling was undertaken using MinKNOW v25.03.9 and Dorado v1.4.0. ONT libraries were prepared with the Rapid sequencing DNA V14 SQK-RBK114.24 barcoding kit according to the manufacturer’s instructions.

### Bioinformatic analyses

Using a combination of short- and long-read data, high-quality closed or close-to-closure genome assemblies were attained for the nine parental strains. Hybrid assemblies were obtained using Unicycler v0.5.1^44^ followed by error correction with Pilon v1.24^45^. SPANDx v4.2^46^ was used to map Illumina reads to the hybrid assemblies, followed by manual error correction of any remaining variants, prior to comparative analyses against the experimental evolution lineages. For our PAO1 (LMG 12228) strain, ATCC 15692 (NCBI accession GCF_001729505.1) was used for hybrid assembly polishing. tBLASTn analysis (web version 2.17.0+ available at: https://blast.ncbi.nlm.nih.gov/Blast.cgi; accessed 04Mar26) was conducted on the PmrB variants using the ‘Microbes’ database, against both the ‘Complete’ (*n*=2,035) and ‘Draft’ (*n*=49,093) *P. aeruginosa* genome databases. This analysis was performed to determine the presence of these variants among a high-quality global genome dataset. Intra-population genomic diversity was analysed using LoFreq v2.1.5^47^. Variants below 5% minor component mixture were excluded from further analysis. AMR determinants were identified using the *P. aeruginosa*-specific database in ARDaP v2.3.2^33^ using default parameters.

## Results

### Phenotypic evolution of high-level CST resistance

Experimental evolution under 14 stepwise, two-fold increasing CST MICs concentrations (i.e. 0.0625, 0.125, 0.25, 0.5, 1, 2, 4, 8, 16, 32, 64, 128, 256, and 512 µg/mL) resulted in visible growth of most lineages up to 512 µg/mL. The exception was SCHI0064.S.1, which failed to grow at 128 µg/mL after 48 hours.

### Convergent evolution and allelic frequency dynamics in *pmrAB* and *phoPQ*

Population-level sequencing was performed across all ten lineages at four CST concentrations (0.5, 4, 64, and 512 µg/mL), except for SCHI0064.S.1 at 512 µg/mL due to extinction prior at >64 μg/mL CST. In total, we identified between 2 to 239 mutations (SNPs and indels) across the adapted lineages compared with their corresponding parental strain. Notably, most (97.4%; range: 92.3-100%) variants were unfixed in the population, remaining as mixed sub-populations, even at high CST concentrations (**Table S1**). The sensor kinase encoded by *pmrB*^48^ was the most frequently mutated gene, with 32 non-synonymous mutations identified, and all adapted lineages encoding at least two *pmrB* mutations (**Table 3**). At 512 μg/mL, seven lineages possessed lineage-specific, dominant (≥50% allele frequency [AF]) PmrB variants: R10L, N16_V19del, V215L, T253M, H261Y, L296Q, and D302N. Several PmrB variants occurred in, or very close to, known hotspot locations (e.g. V9A, R10L and A247T), whereas others were seen in regions not previously reported in CST-resistant strains (**Figure 2**; **Table 3**). Most *pmrB* variants occurred in either the histidine kinase domain (*n*=12/32; 37.5%) or periplasmic loop (*n*=11/32; 34.4%) (**Figure 2**). Some convergence was evident, with four *pmrB* variants (D47del, T132P, L172del and T253M) occurring in more than one lineage (**Table 3**).

We identified three variants (G13S [PAO1], D14G [SCHI0021.S.7 and SCHI0033.S.19] and G15S [SCHI0064.S.1]) in the N-terminal receiver domain of the TCS response regulator, PmrA, which is encoded directly upstream of *pmrB* (**Table 3**). The G13S variant emerged at a relatively low CST concentration (4 μg/mL) and became dominant (∼53.6% AF) at 512 μg/mL (**Table 3**, **Figure 1**). PmrA D14G also emerged at 4 μg/mL and reached 11.8% and 38.4% AFs in SCHI0021.S.7 and SCHI0033.S.19, respectively, by 512 μg/mL CST (**Table 3**, **Figure 1**). PmrA G15S emerged only once, as a minor component (8.2% AF) in the SCHI0064.S.1 lineage at 64 μg/mL.

**Figure 1.**
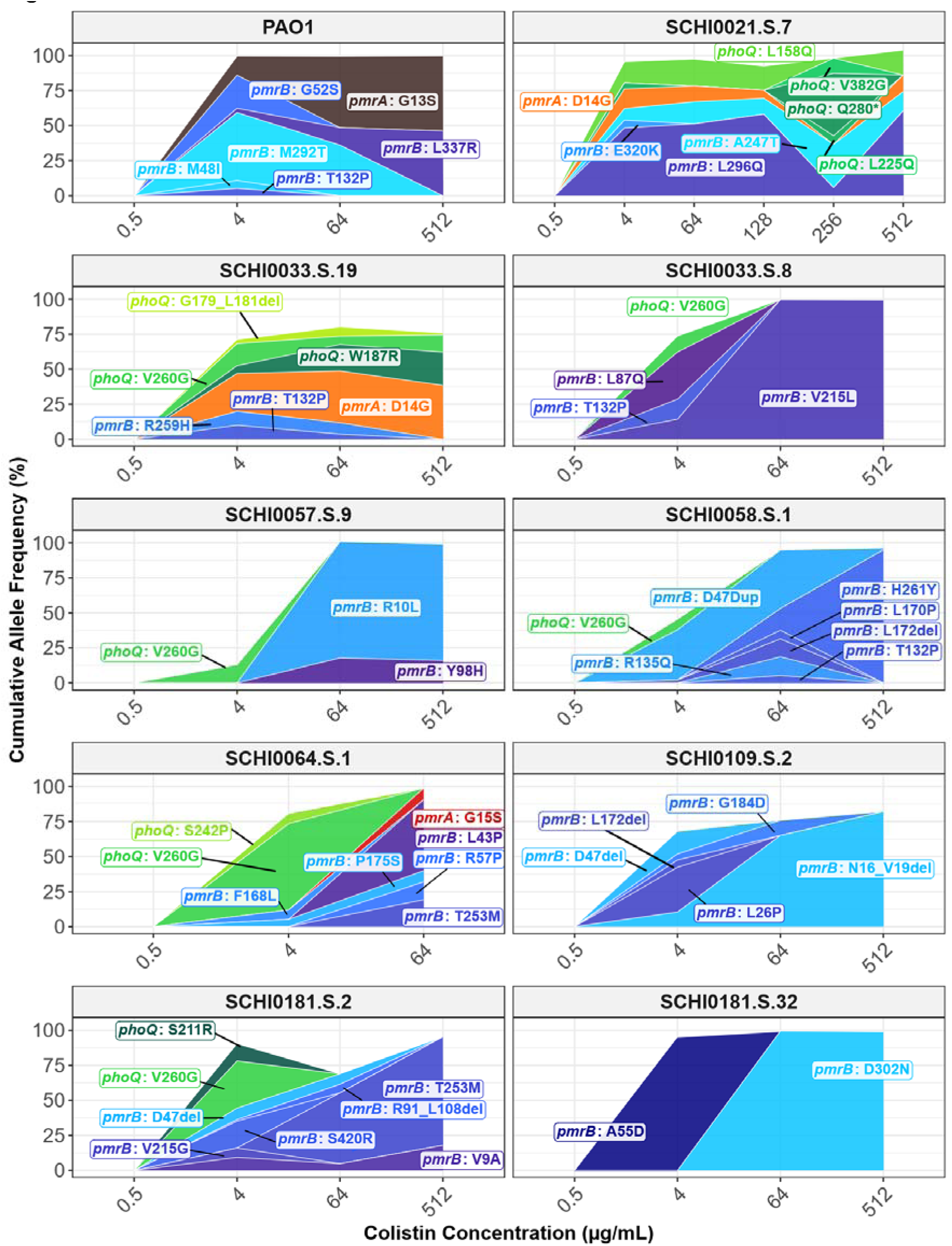
Cumulative mutations across the two-component systems, *pmrAB* and *phoPQ*, and their associated allele frequencies (AFs) during experimental evolution at increasing CST concentrations. Mutations in the sensor kinase gene, *pmrB*, were identified across all CST-resistant lineages whereas mutations in *pmrA* and *phoQ* were rarer. Shading is indicative of the cumulative AF of all known driver mutations occurring in that lineage.

**Figure 2.**
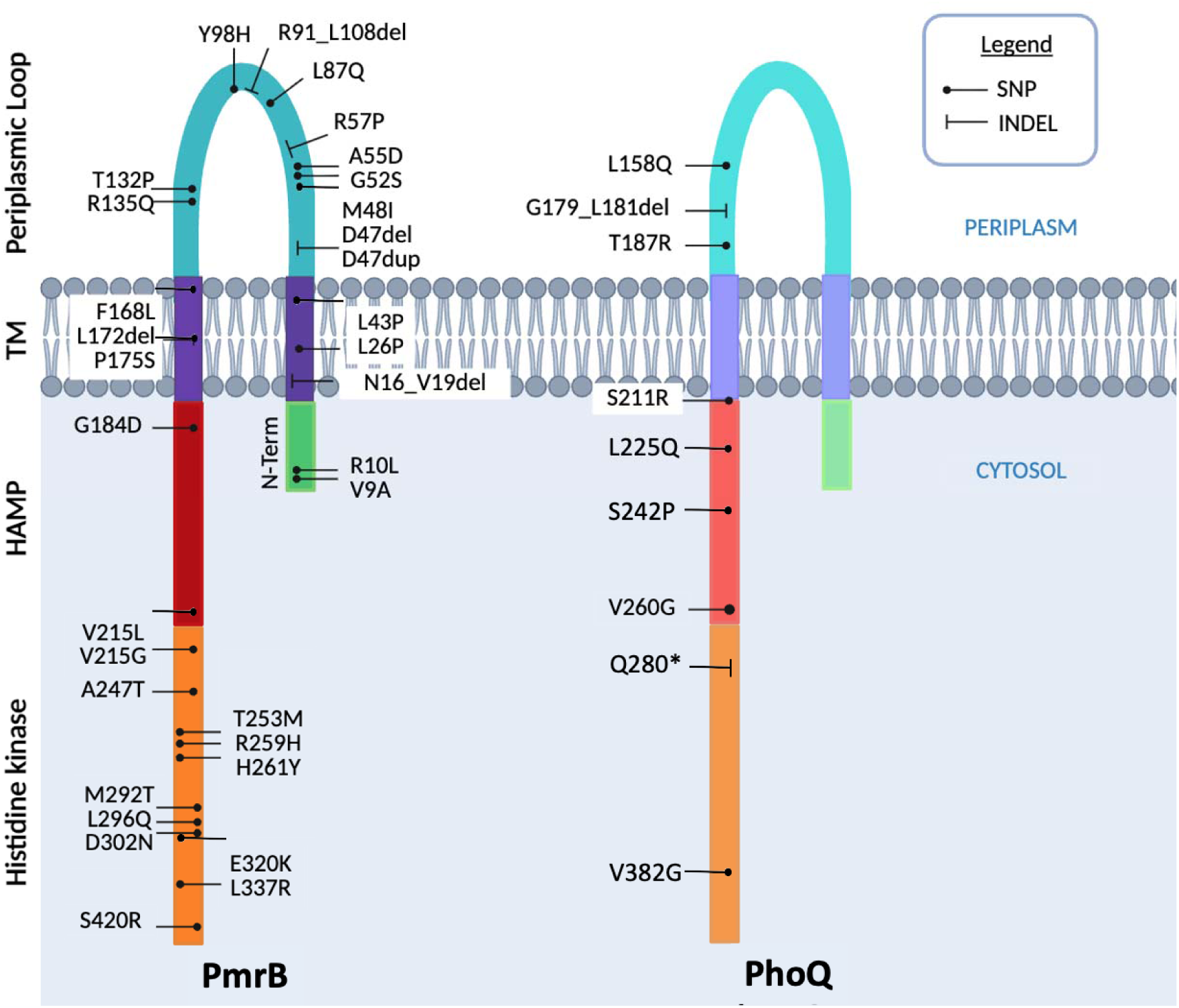
Approximate locations of identified PmrB (left) and PhoQ (right) variants within predicted protein domains. More variants were observed in *pmrB* than *phoQ,* and tended to cluster within either histidine kinase domain or periplasmic loop. Abbreviations: HAMP, histidine kinases, adenyl cyclases, methyl-accepting proteins, and phosphatases; TM, transmembrane; SNP, single nucleotide polymorphism; INDEL, insertion/deletion; *, stop codon.

Although infrequent compared to *pmrAB* mutations, we also observed convergent evolution in another TCS sensor kinase known to play a role in acquired CST resistance, PhoPQ. Seven lineages accumulated *phoQ* mutations, with seven of the nine mutations (L158Q, G179_L181del, T187R, S211R, L225Q, S242P, and Q280stop) not reported previously (**Table 3**; **Figure 1**). PhoQ variants were dominated by a highly convergent hotspot mutation, V260G, which has been frequently observed both *in vitro* and *in vivo*^49^. This variant emerged independently across six lineages (SCHI0033.S.8, SCHI0033.S.19, SCHI0057.S.9, SCHI0058.S.1, SCHI0064.S.1, and SCHI0181.S.2). Notably, *phoQ* mutations never became fixed in any lineage, instead persisting at a minor component (range: 1.3 to 44.7% AF) through increasing selection pressure **(Table 3).** Unlike *pmrAB*, we did not observe any mutations in the PhoPQ TCS response regulator gene, *phoP*.

Whilst canonical mutations within the TCSs *pmrAB* and *phoPQ* accounted for most of the highly resistant populations across the lineages (**Figure 1**), SCHI0033.S.19 and SCHI0109.S.2 presented notable exceptions. In these two populations, the cumulative AFs of primary *pmrAB*/*phoPQ* mutations plateaued at around ∼75-80% at 512 µg/mL. This distinct population vacuum was filled by the emergence of novel secondary driver mutations. In SCHI0033.S.19, the remaining sub-population was largely defined by a unique missense mutation in the biofilm master regulator *amrZ* (S41I; 16.1% AF at 512 µg/mL) and a missense variant in the lipid A modifying gene *lpxO2* (T214P; 10.2% AF at 512 µg/mL). This compensatory dynamic was mirrored in SCHI0109.S.2, where the remaining ∼25% of the population unoccupied by a *pmrB* deletion was filled by a distinct sub-clone harbouring both a deleterious *lpxO2* frameshift (P123fs; 29.2% AF at 512 µg/mL) and a mutation in the outer membrane assembly factor encoded by *opr86* (L510_G511insSVNSL; 33.9% AF at 512 µg/mL).

### Convergent metabolic rewiring drives biofilm formation and outer-membrane remodelling

In addition to canonical TCS mutations, we observed parallel evolution across several regulatory nodes of the cyclic-di-GMP network^50^, likely driving sub-populations toward a sessile, biofilm-forming lifestyle. These mutations generally emerged at low CST concentrations, never established dominance in the population, and were subsequently lost at higher CST levels. Mutations were identified in the *wspF* repressor in SCHI0033.S.8 (G35D, up to 18.0% AF) and SCHI0021.S.7 (Y305stop; 5.7% AF at 0.5 µg/mL), the sensor kinase *wspA* (SCHI0033.S.8, multiple mutations, up to 21.1% AF), and the diguanylate cyclase *wspR_1* (SCHI0109.S.2, multiple mutations, up to 16.4% AF) (Table S1). Some additional mutations were also identified in other cyclic-di-GMP phosphodiesterases and regulators, including a *bifA* truncation in SCHI0021.S.7 (Q206stop) and a *morA* truncation in SCHI0057.S.9 (E1156stop). This regulatory shift was further supported by the emergence of the *amrZ* master regulator variant (S41I; 16.1% AF at 512 µg/mL) in SCHI0033.S.19, and nonsense mutations (H657fs and Trp504stop) in the global *gacS* sensor kinase, which combined to sweep nearly 60% of the SCHI0181.S.32 population at the 512 µg/mL timepoint.

We observed convergent metabolic adaptation in multiple lineages. In SCHI0064.S.1, a missense mutation in glycerol-3-phosphate dehydrogenase (*glpD;* T6A) expanded from an AF of 29.2% at 4 µg/mL to become the dominant allele (77.1% AF) at 64 µg/mL. Additionally, alterations affecting glutamate transport (*gltS* F168L, 15.4% AF in SCHI0021.S.7 at 4 µg/mL) and fructose regulation (*fruR* A54G, 8.5% AF in SCHI0181.S.2 at 512 µg/mL) emerged as secondary minor variants. These metabolic adaptations likely act as compensatory mechanisms to offset the high metabolic burden associated with simultaneous lipid A modification and increased biofilm production.

### Mutator phenotypes and response to oxidative stress

The mutational burden across all lineages was broadly similar; however, SCHI0064.S.1 and SCHI0109.S.2 were notable exceptions (**Figure 3**) due to the emergence of hypermutator phenotypes. Hypermutation in SCHI0064.S.1 was driven by a canonical DNA mismatch repair defect (MutL R264C) that emerged at 4 µg/mL CST (26.6% AF) and quickly became dominant by 64 µg/mL (84.7% AF). This lineage consequently exhibited a substantially elevated transition-to-transversion (Ts/Tv) ratio of 9.3 by the final timepoint (**Table S2**), a hallmark signature of DNA mismatch repair deficiency^51^. Hypermutation in SCHI0109.S.2 was driven by a mutY defect (K343T), emerging briefly as a minor component at 64 µg/mL CST before becoming extinct. In contrast to MutL R264C, the MutY K343 substantially decreased the Ts/Tv ratio, crashing to 0.02 at 64 μg/mL before returning to a normal range (∼1.3-2.5 in *P. aeruginosa*). A third putative hypermutator transiently emerged at 64 µg/mL in the SCHI0058.S.1 lineage (*mutL* A406G, 6.1% AF), with a modestly elevated Ts/Tv ratio (3.9), but became extinct by the next timepoint with no visible shift in the Ts/Tv signatures (**Figure 3; Table S2**). Although no defect in the mismatch repair system was identified in the SCHI0033.S.8, this lineage rapidly accumulated mutations prior to the 0.5 µg/mL timepoint, quickly plateauing for the remainder of the timepoints, suggesting the brief emergence of a hypermutator that became extinct prior to sampling.

**Figure 3.**
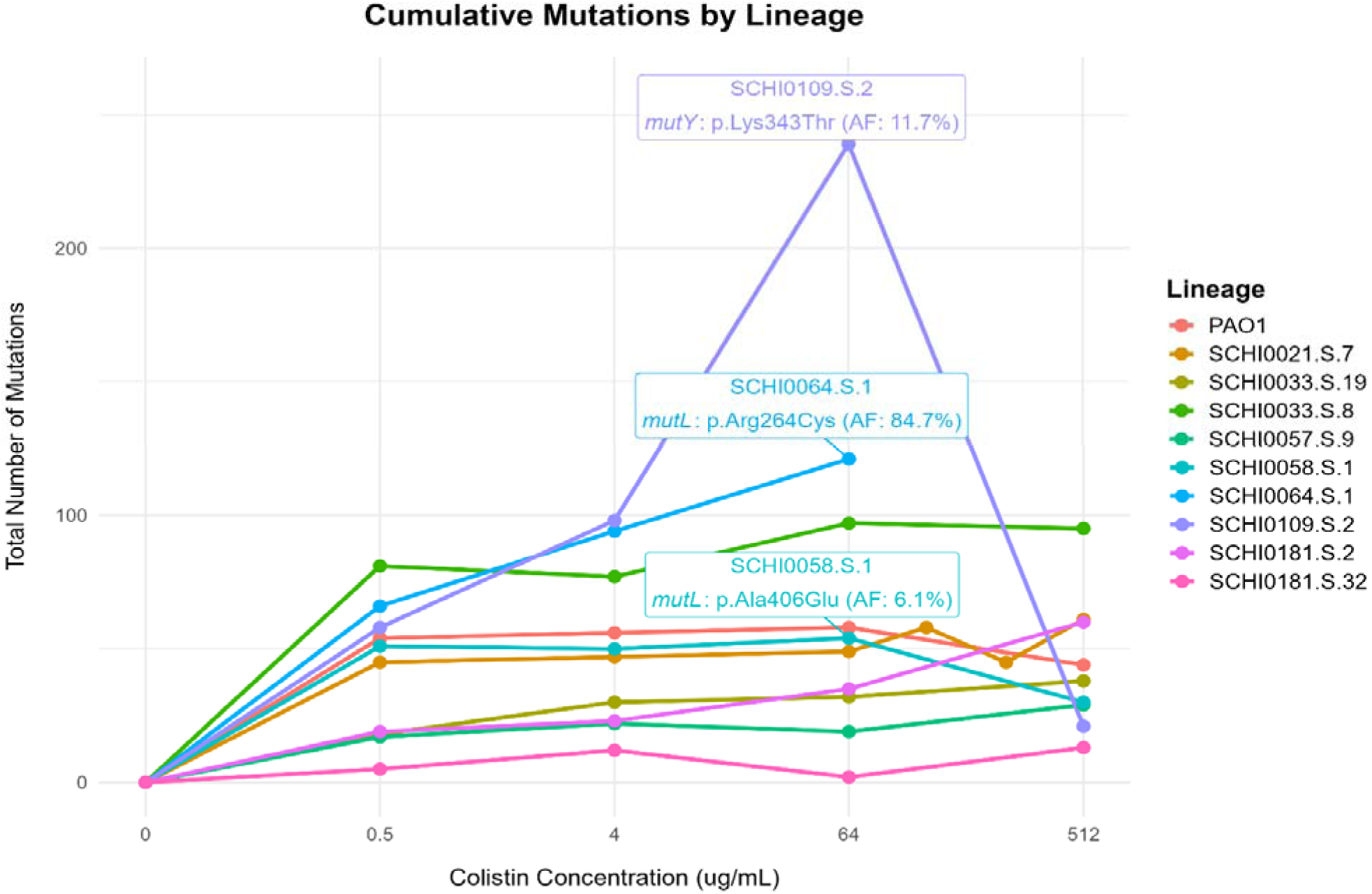
Cumulative mutations across all lineages. Mutations include both single-nucleotide polymorphisms and insertions/deletions. The SCHI0064.S.1 lineage accumulated the most mutations by the 64 _μ_g/mL colistin minimum inhibitory level due to a MutL defect, resulting in a hypermutator phenotype.

In contrast to the systemic hypermutators, most populations exhibited standard mutational profiles with normal or mildly elevated Ts/Tv ratios (range 0.6–5.4; **Table S2**). Despite this overall genomic stability, the acute oxidative stress induced by CST caused multiple compensatory adaptations to manage widespread intracellular damage. For instance, lineage SCHI0181.S.32 developed a missense mutation in the master molecular chaperone *dnaK* (T36P), which rapidly became fixed (98.6% AF at 0.5 µg/mL), highlighting the need to manage oxidised, misfolded proteins before canonical resistance mechanisms fully emerge.

Most lineages evolved convergent, low-frequency sub-clones, likely to compensate for the structural DNA damage caused by reactive oxygen species (ROS). Mutations emerged in the damage-inducible helicase *dinG* (V201delinsAL) and the recombinase *xerD* (H270R) across multiple independent lineages at AFs between 6-11%, likely facilitating DNA repair and increased genomic stability in response to ROS toxicity (**Table S1**). Concurrently, six independent lineages (PAO1, SCHI0021.S.7, SCHI0033.S.8, SCHI0058.S.1, SCHI0109.S.2, and SCHI0181.S.2) acquired a convergent mutation in the transcription-repair coupling factor *mfd* (A445V; ∼5-12% AF) and three independent lineages (SCHI0021.S.7, SCHI0033.S.8, SCHI0064.S.1) developed mutations in the *recA* recombinase (I79V: SCHI0021.S.7; 14.4% AF at 0.5 µg/mL, SCHI0033.S.8; 11.5% AF at 64 µg/mL and SCHI0064.S.1; 8.7% AF at 0.5 µg/mL, and Q83P: SCHI0033.S.8; 11.0% AF at 4 µg/mL).

## Discussion

High-level CST resistance in *P. aeruginosa* is traditionally viewed through the narrow lens of lipid A modification, typically within the setting of laboratory-domesticated PAO1 lineages. However, our experimental evolution of contemporary clinical lineages reveals a far more complex, multi-tiered adaptive strategy for CST resistance development. We demonstrate that survival under escalating CST pressure is not achieved through singular, stepwise mutations, but rather through a dynamic interplay of intense clonal competition, transient hypermutation, two-component system (TCS) mutation, and mutualistic metabolic adaptations. For all strains, we observed a universal accumulation of mutations within the *pmrAB* TCSs, frequently accompanied by *phoQ* mutations, substantiating the role of these two TCSs as primary drivers of CST resistance. Our recovery of canonical mutations, including *pmrB* V9A^52^, R10L^53^, and *phoQ* V260G^28^, aligns with prior evolutionary models and validates the intense selective pressure acting on these TCSs. Despite the known breadth of the *P. aeruginosa* regulatory network, where alternative TCSs such as ParRS, CprRS, and ColRS have been implicated in CST resistance^54^, we did not observe any mutations in these systems. Instead, our study shows that the extensive diversity of novel *pmrB* variants reveals an underappreciated flexibility in how TCS activation can be achieved, which may be further influenced by genetic background. Furthermore, the extent of allelic turnover at intermediate CST concentrations highlights a landscape of clonal interference, likely reflecting a trade-off between increasing CST tolerance versus fitness cost before definitive high-level resistance mutations become dominant in the population.

Several *pmrB* variants identified at high CST concentrations have been reported in phenotypically resistant clinical isolates, validating their role in CST AMR. For example, PmrB R135Q, T253M, and R259H have previously been reported in CST-resistant CF lineages^55^. In our study, we identified these variants in lineages sourced from COPD, bronchiectasis, or bloodstream infections, highlighting that these evolutionary trajectories are not exclusive to CF, but rather demonstrate convergence across diverse disease states and *P. aeruginosa* genetic backgrounds. Interestingly, multiple studies have shown that clinical isolates frequently harbour multiple *pmrB* mutations co-occurring within a single genome (e.g., R10L alongside Y345H, or combinations involving R79H, A95T, and M292I) to achieve high MICs through an additive effect^56^. Although our evolved lineages also contained multiple, simultaneously occurring *pmrB* variants (e.g. R10L coexisting with Y98H in SCHI0057.S.9), our deep sequencing approach aimed to capture population-level dynamics rather than instances of co-occurrence. Therefore, rather than representing compounded mutations within a single genome, the multiple *pmrB* variants identified in our study likely reflects intense clonal interference, where multiple distinct resistant sub-clones continuously compete within the co-culture.

Comparing these population dynamics against both clinical and experimental literature is critical for deciphering mutational noise from true resistance drivers. For instance, we found that PmrB A247T persisted across CST-resistant populations at 64 to 512 µg/mL concentrations in the SCHI0021.S.7 lineage. Interestingly, this variant has been also observed in an experimental microdroplet model^57^ and clinical strains^58,59^, and has been functionally verified to confer CST resistance in genetic manipulation work^60^. This highly conserved region contains the active phosphorylation pocket and serves as the critical structural switch for PmrB, with mutations in this domain frequently mimicking the phosphorylated state, leading to constitutive, kinase-independent activation of the downstream resistance regulon^61,62^.

Although canonical *pmrAB* and *phoPQ* mutations drove resistance in most lineages, our data uncovered alterations in systems other than TCSs can also drive resistance. In lineages SCHI0033.S.19 and SCHI0109.S.2, primary TCS mutations accounted for only ∼75-80% of the bacterial population at the 512 µg/mL timepoint. This distinct population vacuum was filled by the emergence of novel secondary mutations, specifically in the lipid A-modifying enzyme LpxO2, the biofilm master regulator AmrZ, and the outer membrane assembly factor Opr86. Recent experimental evolution studies indicate that loss-of-function mutations in *lpxO2* severely reduce CST membrane binding under specific stress conditions^63^, demonstrating that altering lipid A hydroxylation provides a compensatory evolutionary trajectory when canonical TCS remodelling is insufficient^52^.

The emergence of hypermutator phenotypes is a recognised strategy for rapid adaptation to CST exposure^52^, and was a defining feature of two of our evolved lineages, both of which were derived from COPD parental strains. Although wild-type *P. aeruginosa* typically exhibits a transition bias (Ts/Tv > 2.0)^64^, these two lineages developed distinct mutational signatures reflecting different mechanisms of DNA repair failure. Lineage SCHI0064.S.1 acquired a canonical MutL DNA mismatch repair defect, which drove a massive influx of uncorrected transitions (Ts/Tv > 9.0) and ultimately caused population collapse after 64 µg/mL. Conversely, lineage SCHI0109.S.2 transiently developed a MutY glycosylase defect. Because MutY normally excises 8-oxoguanine lesions generated by ROS, its impairment caused a rapid, substantial spike in the mutation rate (Figure 3) and a severe transversion bias (Ts/Tv of 0.02). The evolutionary collapse of the defective MutL lineage raises the critical question of whether the high prevalence of hypermutators in CF *P. aeruginosa* infections (up to 60% of case)^65^ contributes to the lower-than-expected CST resistance rate observed in this cohort^65^. Isolates lacking an active DNA mismatch repair system may be extremely susceptible to CST-induced oxidative stress, suffering lethal mutation accumulation before successful adaptations can establish in the population. Taken together, our findings highlight the diverse evolutionary strategies used by *P. aeruginosa* to navigate antimicrobial-induced bottlenecks.

To survive acute ROS-induced damage, the remaining non-hypermutator lineages deployed highly convergent, localised protective responses. For instance, the severity of ROS toxicity was evidenced by a massive population sweep at low CST levels in SCHI0181.S.32 involving the master molecular chaperone *dnaK* (T36P, 98.6% at 0.5 µg/mL). DnaK prevents toxic aggregation of ROS-misfolded proteins^66^, suggesting an immediate need to manage oxidised proteins before canonical resistance fully emerged. Similarly, other lineages had low-frequency sub-clones emerge to rescue collapsed replication forks via the damage-inducible helicase *dinG* and the recombinase *xerD*. Concurrently, the mutation frequency decline (*mfd*) protein, a factor within the transcription-coupled repair pathway, was mutated across six independent lineages indicating its important evolutionary role (Table S1). Recent research has redefined this *mfd* system, demonstrating that transcription-coupled repair is a primary driver of endogenous oxidative stress, leading to increased mutagenesis^67^ and accelerated AMR evolution^68^. Our *mfd* variant convergence suggests that this localised increase in mutagenesis provides a short but critical selective advantage to bypass ROS-induced transcriptional errors. Consistent with models of mutator hitchhiking, which have been validated in longitudinal studies of *P. aeruginosa* adaptation within CF lungs^69^, this transient subpopulation allows the lineage to persist through moderate CST stress before being outcompeted by fitter clones without this defect.

Surviving extreme antibiotic pressure requires mitigating the fitness costs associated with lipid A modification. Recent metabolomic studies demonstrate that polymyxin resistance heavily disrupts intracellular carbohydrate flux, imposing severe metabolic stress and energy limitations^70,71^. Rather than modifying central glycolysis directly, our analysis revealed that convergent mutations likely rewire the cyclic-di-GMP (c-di-GMP) network to shift towards a sessile, biofilm-associated lifestyle, coinciding with increased extrapolysaccharide production^72^. Mutations occurred across multiple regulators and sensory nodes, including the Wsp chemosensory system (*wspA*, *wspF*, *wspR_1*), the GacS/GacA TCS, and the BifA and MorA phosphodiesterases. By disrupting these regulators, lineages derepress cyclic-di-GMP production, reducing flagellar motility and redirecting energy toward synthesising protective exopolysaccharide matrices. However, this altered metabolic profile imposes a significant biosynthetic burden, forcing populations to shunt carbon flux away from standard metabolism. This burden was illustrated in SCHI0064.S.1, which underwent a massive metabolic sweep via the acquisition of a missense GlpD (glycerol-3-phosphate dehydrogenase) mutation (T6A; 77.1% AF). Consistent with previous evolutionary models in *P. aeruginosa*^76,77^, we hypothesise that these localised, matrix-producing sub-populations act cooperatively. This shared public good confers collateral CST protection to the wider community, allowing the population to reach a resistant protective threshold without condemning the entire community to the unsustainable metabolic cost of ubiquitous, matrix overproduction^78^.

We acknowledge some study limitations. Although we identified several mutational drivers of CST resistance and corresponding metabolic adaptations, we were unable to phenotypically characterise individual clones, as removal of CST selective pressure risks reversion of unstable adaptive mutations. Additionally, the direct impact of individual mutations (either directly facilitating CST resistance/tolerance or metabolically supporting fitness costs) was not experimentally validated through targeted genetic manipulation (e.g., allelic exchange or gene knockouts) or transcriptomic analyses. Furthermore, a limitation of all *in vitro* models is the inability to fully replicate or capture the complex selective pressures encountered during chronic respiratory infection, including host immune responses, polymicrobial interactions, and spatial heterogeneity. Finally, while population-level sequencing enables detection of low-frequency variants, it cannot definitively resolve epistatic interactions or assign co-occurring mutations to the same genome without complementary long-read or single-cell approaches.

In conclusion, we report several new genetic variants that confer CST resistance in *P. aeruginosa*. Using population-level sequencing, we demonstrate that high-level CST resistance evolution in clinical *P. aeruginosa* isolates involves complex and intricate convergent evolutionary pathways. Furthermore, it is uncommon for a single genetic variant to confer clinically relevant CST resistance (i.e. ≥4 μg/mL), instead requiring compounding mutation accumulation across different genes. We observed a strong parallel initial adaptive response, whereby CST resistance was universally driven by *pmrAB* and *phoPQ* sensor kinase mutations, resulting in lipid A modification. Several lineages gained additional *phoQ* and/or *pmrA* mutations, highlighting the dynamic and complex compounding effect required to reach high-level CST resistance. When primary TCS remodelling was insufficient to survive extreme concentrations, distinct secondary driver mutations (such as those impacting *lpxO2* and *amrZ*) emerged. The severe fitness cost imposed by primary TCS modification, exacerbated by CST-induced metabolic and oxidative stress, resulted in the evolution of numerous metabolic, DNA repair, and membrane modification adaptations, creating heterogenous protective sub-populations. Extreme CST-induced oxidative stress sometimes led to distinct mutational signatures, including a severe transversion bias driven by a MutY repair defect, along with DNA rescue (*dinG*, *xerD*) and localised evolvability factor (*mfd*) defects aimed at altering DNA damage repair pathways. Furthermore, the recurrent acquisition of mutations across the cyclic-di-GMP signalling network (*wspF*, *wspA*, *gacS*, *bifA*, and *morA*) and metabolic shunts (*glpD*) increased biofilm production, possibly protecting the broader community during CST adaptation. Together, our findings demonstrate that high-level CST resistance in clinical *P. aeruginosa* isolates requires a holistic rewiring of the bacterium’s metabolic and stress-response networks alongside canonical TCS mutations.

## Acknowledgements

We are grateful to The Mater Research David Serisier Respiratory Biobank curators (Megan Martin, Lucy Burr, and Simon Bowler) for providing access to retrospective bronchiectasis sputa.

## Data Availability

All data included in the study is publicly available in BioProject PRJNA1513741.

## References

1. World Health Organization. WHO publishes list of bacteria for which new antibiotics are urgently needed. https://www.frontiersin.org/articles/10.3389/fcimb.2022.926758/full#B49 (2017).

2. Global AMR R&D & World Health Organization. Incentivising the Development of New Antibacterial Treatments 2023 Progress Report by the Global AMR R&D Hub & WHO. https://cdn.who.int/media/docs/default-source/antimicrobial-resistance/amr-gcp-irc/incentivising-development-of-new-antibacterial-treatments-2023---progress-report.pdf?sfvrsn#72e4f738_3 (2023).

3. Reynolds, D. & Kollef, M. The Epidemiology and Pathogenesis and Treatment of Pseudomonas aeruginosa Infections: An Update. Drugs 81, 2117–2131 (2021).

4. Bhagirath, A. Y. et al. Cystic fibrosis lung environment and Pseudomonas aeruginosa infection. BMC Pulm. Med. 16, 174 (2016).

5. Finch, S., McDonnell, M. J., Abo-Leyah, H., Aliberti, S. & Chalmers, J. D. A Comprehensive Analysis of the Impact of *Pseudomonas aeruginosa* Colonisation on Prognosis in Adult Bronchiectasis. Ann. Am. Thorac. Soc. AnnalsATS.201506–333OC (2015) doi:10.1513/AnnalsATS.201506-333OC.

6. Li, J. et al. Colistin: the re-emerging antibiotic for multidrug-resistant Gram-negative bacterial infections. Lancet Infect. Dis. 6, 589–601 (2006).

7. Narimisa, N. et al. Prevalence of colistin resistance in clinical isolates of Pseudomonas aeruginosa: a systematic review and meta-analysis. Front. Microbiol. 15, 1477836 (2024).

8. O’Carroll, M. R. et al. Clonal strains of Pseudomonas aeruginosa in paediatric and adult cystic fibrosis units. Eur. Respir. J. 24, 101–106 (2004).

9. Damtie, M. A., Vijay, A. K. & Willcox, M. D. P. Detection of Genes Associated with Polymyxin and Antimicrobial Peptide Resistance in Isolates of Pseudomonas aeruginosa. Int. J. Mol. Sci. 26, 10499 (2025).

10. Strickland, K. R., Jelocnik, M., Price, E. P. & Sarovich, D. S. Prevalence of Pseudomonas aeruginosa in Australian wild birds, native wildlife, livestock and domestic animals. Sci. Rep. 16, 15423 (2026).

11. Gogry, F. A., Siddiqui, M. T., Sultan, I. & Haq, Q. Mohd. R. Current Update on Intrinsic and Acquired Colistin Resistance Mechanisms in Bacteria. Front. Med. 8, 677720 (2021).

12. Olaitan, A. O., Morand, S. & Rolain, J.-M. Mechanisms of polymyxin resistance: acquired and intrinsic resistance in bacteria. Front. Microbiol. 5, (2014).

13. Vatansever, C. et al. Co-existence of OXA-48 and NDM-1 in colistin resistant *Pseudomonas aeruginosa* ST235. Emerg. Microbes Infect. 9, 152–154 (2020).

14. Loi, D. V. et al. Clinical Outcomes of Aerosolized Versus Intravenous Colistin in Ventilator-Associated Pneumonia Caused by Multidrug-Resistant Gram-Negative Bacteria. J. Clin. Med. Res. 18, 42–49 (2026).

15. Karaiskos, I. et al. High-Dose Nebulized Colistin Methanesulfonate and the Role in Hospital-Acquired Pneumonia Caused by Gram-Negative Bacteria with Difficult-to-Treat Resistance: A Review. Microorganisms 11, 1459 (2023).

16. De Pascale, G. et al. Use of High-Dose Nebulized Colistimethate in Patients with Colistin-Only Susceptible Acinetobacter baumannii VAP: Clinical, Pharmacokinetic and Microbiome Features. Antibiotics 12, 125 (2023).

17. Davoodi, N. R., Soleimani, N., Hosseini, S. M. & Rahnamaye-Farzami, M. Co-occurrence of mcr-1 and mcr-3 mobilized colistin resistance genes among carbapenem-resistant Pseudomonas aeruginosa in Iran. Sci. Rep. 16, 833 (2025).

18 . Abd El-Baky, R. M., et al. Prevalence and Some Possible Mechanisms of Colistin Resistance Among Multidrug-Resistant and Extensively Drug-Resistant Pseudomonas aeruginosa. Infect. Drug Resist. 13, 323–332 (2020).

19. Hameed, F. et al. Plasmid-mediated mcr-1 gene in Acinetobacter baumannii and Pseudomonas aeruginosa: first report from Pakistan. Rev. Soc. Bras. Med. Trop. 52, e20190237 (2019).

20. Bialvaei, A. Z. & Samadi Kafil, H. Colistin, mechanisms and prevalence of resistance. Curr. Med. Res. Opin. 31, 707–721 (2015).

21. Andrade, F. F., Silva, D., Rodrigues, A. & Pina-Vaz, C. Colistin Update on Its Mechanism of Action and Resistance, Present and Future Challenges. Microorganisms 8, 1716 (2020).

22. Tran, T. B. et al. Pharmacokinetics/pharmacodynamics of colistin and polymyxin B: are we there yet? Int. J. Antimicrob. Agents 48, 592–597 (2016).

23. El-Sayed Ahmed, M. A. E.-G., et al. Colistin and its role in the Era of antibiotic resistance: an extended review (2000–2019). Emerg. Microbes Infect. 9, 868– 885 (2020).

24. Gurjar, M. Colistin for lung infection: an update. J. Intensive Care 3, 3 (2015).

25. Lim, L. M. et al. Resurgence of Colistin: A Review of Resistance, Toxicity, Pharmacodynamics, and Dosing. Pharmacother. J. Hum. Pharmacol. Drug Ther. 30, 1279–1291 (2010).

26. Sharma, J., Sharma, D., Singh, A. & Sunita, K. Colistin Resistance and Management of Drug Resistant Infections. Can. J. Infect. Dis. Med. Microbiol. 2022, 1–10 (2022).

27. Nang, S. C., Azad, M. A. K., Velkov, T., Zhou, Q. (Tony) & Li, J. Rescuing the Last-Line Polymyxins: Achievements and Challenges. Pharmacol. Rev. 73, 679– 728 (2021).

28. Jochumsen, N. et al. The evolution of antimicrobial peptide resistance in Pseudomonas aeruginosa is shaped by strong epistatic interactions. Nat. Commun. 7, 13002 (2016).

29. Disney-McKeethen, S., Seo, S., Mehta, H., Ghosh, K. & Shamoo, Y. Experimental evolution of Pseudomonas aeruginosa to colistin in spatially confined microdroplets identifies evolutionary trajectories consistent with adaptation in microaerobic lung environments. mBio 14, e01506–23.

30. Alarcon Rios, A. C., et al. Colistin resistance dynamics in Pseudomonas aeruginosa under biofilm and planktonic growth. Antimicrob. Agents Chemother. 69, e00421–25.

31. Cervoni, M., et al. The Genetic Background and Culture Medium Only Marginally Affect the In Vitro Evolution of Pseudomonas aeruginosa Toward Colistin Resistance. Antibiotics 14, (2025).

32. Hsieh, Y.-Y. P. et al. Magnesium depletion unleashes two unusual modes of colistin resistance with different fitness costs. BioRxiv Prepr. Serv. Biol. 2024.10.15.618514 (2025) doi:10.1101/2024.10.15.618514.

33. Madden, D. E. et al. Keeping up with the pathogens: improved antimicrobial resistance detection and prediction from Pseudomonas aeruginosa genomes. Genome Med. 16, 78 (2024).

34. Parkins, M. D., Somayaji, R. & Waters, V. J. Epidemiology, Biology, and Impact of Clonal Pseudomonas aeruginosa Infections in Cystic Fibrosis. Clin. Microbiol. Rev. 31, 10.1128/cmr.00019-18 (2018).

35. Webb, K. A. et al. Genomic diversity and antimicrobial resistance of Prevotella species isolated from chronic lung disease airways. *Microb*. Genomics 8, 000754 (2022).

36. Stewart, A. G. et al. Molecular Epidemiology of Third-Generation-Cephalosporin-Resistant Enterobacteriaceae in Southeast Queensland, Australia. Antimicrob. Agents Chemother. 65, e00130–21 (2021).

37. McCarthy, K. L. & Paterson, D. L. Long-term mortality following Pseudomonas aeruginosa bloodstream infection. J. Hosp. Infect. 95, 292–299 (2017).

38. Chichón, G. et al. Spread of Pseudomonas aeruginosa ST274 Clone in Different Niches: Resistome, Virulome, and Phylogenetic Relationship. Antibiotics 12, 1561 (2023).

39. Holloway, B. W. Genetic Recombination in Pseudomonas aeruginosa. Microbiology 13, 572–581 (1955).

40. Madden, D. E., et al. Express Yourself: Quantitative Real-Time PCR Assays for Rapid Chromosomal Antimicrobial Resistance Detection in Pseudomonas aeruginosa. Antimicrob. Agents Chemother. 66, e0020422 (2022).

41. Madden, D. E. et al. Express Yourself: Quantitative Real-Time PCR Assays for Rapid Chromosomal Antimicrobial Resistance Detection in Pseudomonas aeruginosa. Antimicrob. Agents Chemother. 66, e00204–22 (2022).

42. Karvanen, M., Malmberg, C., Lagerbäck, P., Friberg, L. E. & Carsp, O. Colistin Is Extensively Lost during Standard *In Vitro* Experimental Conditions. Antimicrob. Agents Chemother. 61, e00857–17 (2017).

43. Li, J., Milne, R. W., Nation, R. L., Turnidge, J. D. & Coulthard, K. Stability of Colistin and Colistin Methanesulfonate in Aqueous Media and Plasma as Determined by High-Performance Liquid Chromatography. Antimicrob. Agents Chemother. 47, 1364–1370 (2003).

44. Wick, R. R., Judd, L. M., Gorrie, C. L. & Holt, K. E. Unicycler: Resolving bacterial genome assemblies from short and long sequencing reads. PLOS Comput. Biol. 13, e1005595 (2017).

45. Walker, B. J. et al. Pilon: An Integrated Tool for Comprehensive Microbial Variant Detection and Genome Assembly Improvement. PLOS ONE 9, e112963 (2014).

46. Sarovich, D. S. & Price, E. P. SPANDx: a genomics pipeline for comparative analysis of large haploid whole genome re-sequencing datasets. BMC Res. Notes 7, 618 (2014).

47. Wilm, A. et al. LoFreq: a sequence-quality aware, ultra-sensitive variant caller for uncovering cell-population heterogeneity from high-throughput sequencing datasets. Nucleic Acids Res. 40, 11189–11201 (2012).

48. Moskowitz, S. M., Ernst, R. K. & Miller, S. I. PmrAB, a two-component regulatory system of Pseudomonas aeruginosa that modulates resistance to cationic antimicrobial peptides and addition of aminoarabinose to lipid A. J. Bacteriol. 186, 575–579 (2004).

49. Barrow, K. & Kwon, D. H. Alterations in Two-Component Regulatory Systems of phoPQ and pmrAB Are Associated with Polymyxin B Resistance in Clinical Isolates of Pseudomonas aeruginosa. Antimicrob. Agents Chemother. 53, 5150– 5154 (2009).

50. Güvener, Z. T. & Harwood, C. S. Subcellular location characteristics of the Pseudomonas aeruginosa GGDEF protein, WspR, indicate that it produces cyclic-di-GMP in response to growth on surfaces. Mol. Microbiol. 66, 1459–1473 (2007).

51. Weigand, M. R. & Sundin, G. W. General and inducible hypermutation facilitate parallel adaptation in Pseudomonas aeruginosa despite divergent mutation spectra. Proc. Natl. Acad. Sci. U. S. A. 109, 13680–13685 (2012).

52. Dößelmann, B. et al. Rapid and Consistent Evolution of Colistin Resistance in Extensively Drug-Resistant Pseudomonas aeruginosa during Morbidostat Culture. Antimicrob. Agents Chemother. 61, 10.1128/aac.00043-17 (2017).

53. Oliver, A., Cantón, R., Campo, P., Baquero, F. & Blázquez, J. High Frequency of Hypermutable Pseudomonas aeruginosa in Cystic Fibrosis Lung Infection. Science 288, 1251–1253 (2000).

54. Fernández, L. et al. Adaptive resistance to the ‘last hope’ antibiotics polymyxin B and colistin in Pseudomonas aeruginosa is mediated by the novel two-component regulatory system ParR-ParS. Antimicrob. Agents Chemother. 54, 3372–3382 (2010).

55. Moskowitz, S. M. et al. PmrB Mutations Promote Polymyxin Resistance of Pseudomonas aeruginosa Isolated from Colistin-Treated Cystic Fibrosis Patients. Antimicrob. Agents Chemother. 56, 1019–1030 (2012).

56. Rossitto, M. et al. The Challenging Life of Mutators: How Pseudomonas aeruginosa Survives between Persistence and Evolution in Cystic Fibrosis Lung. Microorganisms 12, 2051 (2024).

57. Disney-McKeethen, S., Seo, S., Mehta, H., Ghosh, K. & Shamoo, Y. Experimental evolution of Pseudomonas aeruginosa to colistin in spatially confined microdroplets identifies evolutionary trajectories consistent with adaptation in microaerobic lung environments. mBio 14, e01506–23 (2023).

58. De Sousa, T. et al. Mutational Analysis of Colistin-Resistant Pseudomonas aeruginosa Isolates: From Genomic Background to Antibiotic Resistance. Pathogens 14, 387 (2025).

59. Fisher, L. W. S., Thorpe, H. A., Sassera, D., Corander, J. & Bryant, J. M. High frequency body site translocation of nosocomial Pseudomonas aeruginosa. Nat. Commun. 16, 9862 (2025).

60. Owusu-Anim, D. & Kwon, D. H. Differential Role of Two-Component Regulatory Systems (phoPQ and pmrAB) in Polymyxin B Susceptibility of Pseudomonas aeruginosa. Adv. Microbiol. 2, (2012).

61. Bourret, R. B. Receiver domain structure and function in response regulator proteins. Curr. Opin. Microbiol. 13, 142–149 (2010).

62. Lee, J.-Y. & Ko, K. S. Mutations and expression of PmrAB and PhoPQ related with colistin resistance in *Pseudomonas aeruginosa* clinical isolates. Diagn. Microbiol. Infect. Dis. 78, 271–276 (2014).

63. Hsieh, Y.-Y. P. et al. Magnesium depletion by Candida albicans unleashes two unusual modes of colistin resistance in Pseudomonas aeruginosa with different fitness costs. PLOS Biol. 24, e3003673 (2026).

64. Spencer, D. H. et al. Whole-Genome Sequence Variation among Multiple Isolates of Pseudomonas aeruginosa. J. Bacteriol. 185, 1316–1325 (2003).

65. Ciofu, O., Riis, B., Pressler, T., Poulsen, H. E. & Høiby, N. Occurrence of hypermutable Pseudomonas aeruginosa in cystic fibrosis patients is associated with the oxidative stress caused by chronic lung inflammation. Antimicrob. Agents Chemother. 49, 2276–2282 (2005).

66. da Cruz Nizer, W. S., et al. Oxidative Stress Response in Pseudomonas aeruginosa. Pathogens 10, 1187 (2021).

67. Oxidative stress drives mutagenesis through transcription-coupled repair in bacteria. https://www.pnas.org/doi/10.1073/pnas.2300761120 doi:10.1073/pnas.2300761120.

68. Carvajal-Garcia, J., et al. A small molecule that inhibits the evolution of antibiotic resistance. NAR Mol. Med. 1, ugae001 (2024).

69. Rossitto, M. et al. The Challenging Life of Mutators: How Pseudomonas aeruginosa Survives between Persistence and Evolution in Cystic Fibrosis Lung. Microorganisms 12, 2051 (2024).

70. Hsieh, Y.-Y. P. et al. Magnesium depletion unleashes two unusual modes of colistin resistance with different fitness costs. BioRxiv Prepr. Serv. Biol. 2024.10.15.618514 (2025) doi:10.1101/2024.10.15.618514.

71. Han, M.-L. et al. Alterations of Metabolic and Lipid Profiles in Polymyxin-Resistant Pseudomonas aeruginosa. Antimicrob. Agents Chemother. 62, e02656–17 (2018).

72. Gloag, E. S. et al. Pseudomonas aeruginosa biofilm-deficient mutants undergo parallel evolution during chronic infection. J. Bacteriol. 208, e00520–25 (2026).

73. Xavier, J. B., Kim, W. & Foster, K. R. A molecular mechanism that stabilizes cooperative secretions in Pseudomonas aeruginosa. Mol. Microbiol. 79, 166–179 (2011).

74. Mellbye, B. & Schuster, M. Physiological framework for the regulation of quorum sensing-dependent public goods in Pseudomonas aeruginosa. J. Bacteriol. 196, 1155–1164 (2014).

75. Billings, N. et al. The Extracellular Matrix Component Psl Provides Fast-Acting Antibiotic Defense in Pseudomonas aeruginosa Biofilms. PLoS Pathog. 9, e1003526 (2013).

76. Mehta, H. H., Prater, A. G. & Shamoo, Y. Using experimental evolution to identify druggable targets that could inhibit the evolution of antimicrobial resistance. J. Antibiot. (Tokyo) 71, 279–286 (2018).

77. Ghassani, A. et al. Mutations in genes lpxL1, bamA, and pmrB impair the susceptibility of cystic fibrosis strains of Pseudomonas aeruginosa to murepavadin. Antimicrob. Agents Chemother. 68, e0129823 (2024).

78. Roson-Calero, N. et al. In vitro potentiation of tetracyclines in Pseudomonas aeruginosa by RW01, a new cyclic peptide. Antimicrob. Agents Chemother. 69, e0145924 (2025).

79. Romano, K. P. et al. Mutations in pmrB Confer Cross-Resistance between the LptD Inhibitor POL7080 and Colistin in Pseudomonas aeruginosa. Antimicrob. Agents Chemother. 63, e00511–19 (2019).

80. Romano, K. P. et al. Mutations in pmrB Confer Cross-Resistance between the LptD Inhibitor POL7080 and Colistin in Pseudomonas aeruginosa. Antimicrob. Agents Chemother. 63, e00511–19 (2019).

81. Bolard, A. et al. Production of Norspermidine Contributes to Aminoglycoside Resistance in pmrAB Mutants of Pseudomonas aeruginosa. Antimicrob. Agents Chemother. 63, e01044–19 (2019).

82. Grall, N. et al. No Emergence of Colistin Resistance in the Respiratory Tract of Lung Transplant Patients Treated With Inhaled Colistin. Transpl. Int. 37, 13545 (2025).

83. Abraham, N. & Kwon, D. H. A single amino acid substitution in PmrB is associated with polymyxin B resistance in clinical isolate of Pseudomonas aeruginosa. FEMS Microbiol. Lett. 298, 249–254 (2009).

84. Schniederjans, M., Koska, M. & Häussler, S. Transcriptional and Mutational Profiling of an Aminoglycoside-Resistant Pseudomonas aeruginosa Small-Colony Variant. Antimicrob. Agents Chemother. 61, e01178–17 (2017).

85. Chebotar, I. et al. Genetic Alternatives for Experimental Adaptation to Colistin in Three Pseudomonas aeruginosa Lineages. Antibiotics 13, (2024).

86. Erdmann, M. B., Gardner, P. P. & Lamont, I. L. The PitA protein contributes to colistin susceptibility in Pseudomonas aeruginosa. PloS One 18, e0292818 (2023).

87. Xiao, Y., Li, K., Zhao, R., Wang, Z. & Liu, W. Molecular characteristics of chromosome-mediated colistin resistance in carbapenem-resistant Pseudomonas aeruginosa isolates from a tertiary hospital in China. BMC Microbiol. 26, 313 (2026).

88. Lin, J. et al. Resistance and Heteroresistance to Colistin in Pseudomonas aeruginosa Isolates from Wenzhou, China. Antimicrob. Agents Chemother. 63, 10.1128/aac.00556-19 (2019).

89. Lindon, S. et al. Antibiotic resistance alters the ability of Pseudomonas aeruginosa to invade bacteria from the respiratory microbiome. Evol. Lett. 8, 735–747 (2024).

